# Biodiversity Of Bacteriophages From Lysed Batch Culture Of Recombinant *Escherichia Coli* BL21(DE3) Examined By Anion Exchange Chromatography And TEM

**DOI:** 10.1101/2020.01.15.907352

**Authors:** Alla Kushkina, Fedor Tovkach

## Abstract

It was found that the phage population assaulted a batch culture of an industrial recombinant derivative of *E. coli* BL21(DE3) was attenuated that manifested in producing pinprick-type plaques and their inability to propagation in subsequent passages. Because of this, the goals of the present research were to evaluate biodiversity and scrutinize the possible virion structural defects of the attenuated phage population prior to pure phage lines isolation. The anion exchange chromatography (AEC) was chosen as the principal method allowing to get a comparative phage population profile based on the virion net surface charge, as well as to treat the 5-liter virion-contained sample and collect high-quality concentrated and separated phage fractions to further analysis. The isolate consisted of a mix of two phages belonging to *Myoviridae* (A2-morphotype) and *Siphoviridae* (B1-morphotype) families. By restriction analysis, the main portion of this phage pool (about 99% of all virions) was identified as the primary population of myophage Lw1, which possessed its own intra-population biodiversity (heterogeneity). It consisted of a major and two minor subpopulations that differed by phage capsid size and shape. The subpopulation III consisted of aberrant tubby-phages with triprolate (expanded at both sides) capsids having low aspect ratio.

## 1 Introduction

Sporadic phage infections (phage lysis) of biotechnological bacterial cultures are omnipresent and lead to heavy financial losses. Despite the fact that the *E. coli* BL21(DE3) system for recombinant genes expression [1] is world widely used in both industry and academics, there are only two reports on phages infecting this bacterium and its derivatives. In 2010 Li et al. reported the complete genome sequence of a *Siphoviridae* family phage that had assaulted an engineered *E. coli* strain [2]. In 2013 we analyzed a complete genome sequence of lytic phage Lw1 of *Myoviridae* family that had caused sporadic lysis of *E. coli*-based producer of human insulin [3]. Investigating this case, we have observed the phage lysate containing about 10^10^–10^11^ PFU /ml that formed plaques on a lawn of parental *E. coli* host under the industrial growth temperature (37 °C). Moreover, most of the appeared plaques failed to propagate in subsequent passages. Because of this, we analyzed the bioreactor phage isolate (the primary Lw1 phage population) looking for virion structural aberrations that might be associated with the attenuation of the virus population, as well as the phage population structure and its biodiversity in general (ratio of normal virions to deviated phage virions). Anion exchange chromatography (AEC) was chosen as the principal method for purification and separation of virions. This made it possible to evaluate the biodiversity of the primary phage isolate and physically segregate this phage pool populations and subpopulations based on a density of virion net surface charge before the phage pure lines isolation step.

## 2 Materials and methods

### 2.1 Sources of phages and bacterial strains

A 5 liter sample of a lysed batch-culture of a recombinant derivative of *E. coli* BL21(DE3) was received directly from a bio-reactor of an industrial biotechnological facility located in Ukraine. The same recombinant *E. coli* strain was used for a plaque assay. The host bacterium bears a recombinant plasmid with human insulin gene.

Phages T4 wild type and Lw1 were from our collection. Strains *E. coli* B^E^ and BL21(DE3) were kindly provided by J.M. Karam (Tulane University, USA), *E. coli* BL21 was purchased at NovaGene.

### 2.2 Bacterial growth medium and phage buffer

LB medium was used in biological assays. Kanamycin 30 µg/ml was added to maintain the recombinant plasmid in the industrial host strain. STM buffer comprising 0.01 M Tris-HCl, pH 7.5, 0.01 M MgSO_4_, 0.05 M NaCl was used for all the phage manipulations.

### 2.3 Phage plaque assay

was conducted by double-layer technique [4]. It was important to use richly filled fresh LB-plates. Test plates were incubated overnight under 37°C.

### 2.4 Phage concentration, purification and separation by DEAE-cellulose column

#### I. Crude phage lysate

was treated with DNAse I (up to 2 U/ml) and chlorophorm at 25 °C, than cell debris was sedimented at 5000 g.

#### II. Anion-exchange chromatography

Fibrous DEAE-cellulose (type 23SS, SERVAGEL, capacity 0.74) in 0.01 M NaP buffer (pH 7) was used according to [5]. NaP buffer 1 M stock solution was made of 1 M Na_2_HPO4 – 57.7 ml, 1 M NaH_2_PO_4_ – 42.3 ml [6]. 0.02– 0.04% sodium azide was applied as conservative. The procedure was carried at 18 °C, natural flow with running speed started up from 15 ml/min. Elution buffers containing 0.15, 0.25 and 0.4 M NaCl were used to fractionate phage particles. Peaks were detected at 260 and 280A spectrofotometrically.

### 2.5 Electrophoresis of native chromatographic fractions

was conducted in 0.8-1% agarose gels in TP buffer 1x (20x stock solution, g/l: Tris – 87, NaH_2_PO4 – 94, Na_2_EDTA – 7.44; pH 8.3) [6] for 2-18 hours, 5-6 V/cm, 120 mA according to principles stated in [7]. DNA bands were detected after ethidium bromide staining by GelAnalyzer image analysis software (http://www.gelanalyzer.com/index.html). The intensity of the DNA band fluorescence was accepted as those one corresponding to the phage concentration, and their values were used to build the fluorescence intensity curves.

### 2.6 CsCl-gradient centrifugation

of the peak fractions was performed in Beckman SW50 rotor (100 000 g, 4 °C, 4h) for additional phage purification and separation. The gradient steps had densities of 1.4, 1.5 and 1.6 g/cm^3^ of CsCl in STM buffer. CsCl-samples were stored at 4 °C and dialyzed against STM buffer when required.

### 2.7 Restriction analysis of total phage DNA

Phage DNA was extracted by the phenol-chloroform method [6]. The samples were analyzed by restriction endonucleases BamHI, BglI, BglII, DpnI, DraI, EcoRI, EcoRV, HindIII, HpaI, MboI, NcoI, NdeI, NotI, PstI, SalI, Sau3A, SmaI, TagI, XhoI manufactured by Fermentas and SybEnzyme accordingly to producer recommendations.

### 2.8 Transverse alternating field electrophoresis (TAFE)

#### I. Extraction of total phage DNA for TAFE

It is known, that phenol treatment can result in fragmentation of DNA [8]. From the other side, dialysis of salt-containing phage samples leads to their diluting and, thus, decreasing of phage particle concentration. To avoid these problems, the phage virion nucleic acid was extracted from CsCl-purified non-dialysed phage samples by osmotic shock and gentle heat lysis with 1% sarcosine in liquid condition. The reaction mix of general volume 40 μl consisted of 1-5 μl of CsCl-containing non-dyalyzed phage sample, sarcosine up to final concentration 1% and TAE buffer up to 0.5x (TEA buffer 20x stock solution, g/l: Tris – 87, Na2EDTA – 7.44, pH 8.0) [6]. CsCl-phage aliquot was diluted immediately with TEA-sarcosine buffer mix to provide osmotic shock conditions that disrupt phage head and lead to virion nucleic acid release with its subsequent conversion into DNA-Cs salt. Then, the mix was heated in a water bath at 60 °C, 10 min. The obtained samples were mixed with standard DNA loading buffer and stored at –20 °C until use.

#### II. TAFE procedure

The DNA-sample loading volume was 10 μl. An optimal DNA concentration in samples was reached by their diluting by 0.5x TAE buffer. DNA samples were separated in TAFE GeneLine Beckman system in 1% agarose gels, 0.5x TAE buffer. Since the analyzed DNA samples were liquid, the gel separation was conducted in two stages: 1) in a horizontal electrophoresis camera – to detention the liquid DNA-samples in gel wells (1 h, 120 mA, 6 V/cm, 25 °C); 2) in a vertical TAFE-camera – the same gel with the detained DNA samples was run with pulse of 9-66 sec (24 h, 120 mA, 9 °C). DNA bands in gels were stained with ethidium bromide. Similar two-step TAFE separation of liquid DNA samples was described in [8]. Marker virion DNAs of phages T4, T5 and T7 were extracted in the same way.

### 2.9 Electron microscopy

Phage samples were adsorbed on copper grids with supporting collodion film strengthened with carbon, stained with 2% uranyl acetate and examined with TEM JEOL 1400 (80 kV, magnification x 20-80 k).

### 2.10 Statistical analysis of the electron microscopy data

Measurements of virions on digital micrographs were performed by ImageJ 1.x software [9]. Each parameter of a virion was presented as a data sample consisting of at least 100 measures (see Supporting General statistic data sheet). Each sample was evaluated by percentiles, restricted if needed and divided into classes by Sturges’s formula [10]:

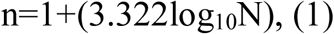

where n – number of classes, N – number of data in a sample. The class interval was calculated by the formula:

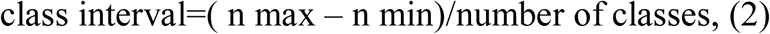

The interval means were used to build a frequency diagram by which the modal value of the parameter analyzed was found.

### 2.11 Calculation of conditional surface area and conditional volume for capsids of *Myovirida*e virions via a model of a prolate spheroid of revolution

It was supposed that elongated icosahedron, which is a shape of the capsid of A2-morphotype phages of *Myoviridae* family (typical representative of this phage morphotype is phage T4 of *E. coli*), could be inscribed within a prolate spheroid of revolution (ellipsoid). Thus, to demonstrate differences in sizes and shapes of myovirion capsids separated into three chromatographic subpopulations, the conditional surface area of capsid was calculated for each virion individually using the formula of the area of ellipsoid:

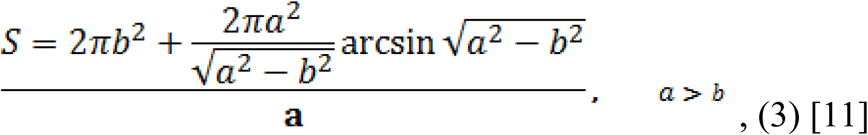

where a - ½ of myophage capsid height, b – ½ of myophage capsid width.

The conditional volume of myophage capsids was estimated by formula for the volume of ellipsoid:

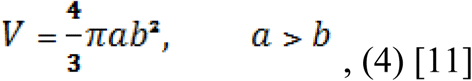

where a and b – the same as in formula (1).

Then, all obtained numerical data were statistically analyzed as described above.

## 3 Results

### 3.1 Plaque morphology

Normally, lytic phages, especially those ones, which are suspected for high virulence, should produce the robust negative colonies on the lawn of the known sensitive host. The phages from the studied primary isolate produced pinprick-type plaques that were of tiny size and poorly differentiated (Fig 1). Most of these negative colonies did not propagate in subsequent passages, but in higher dilutions they completely lysed host cells in a lawn.

**FIGURE 1.**
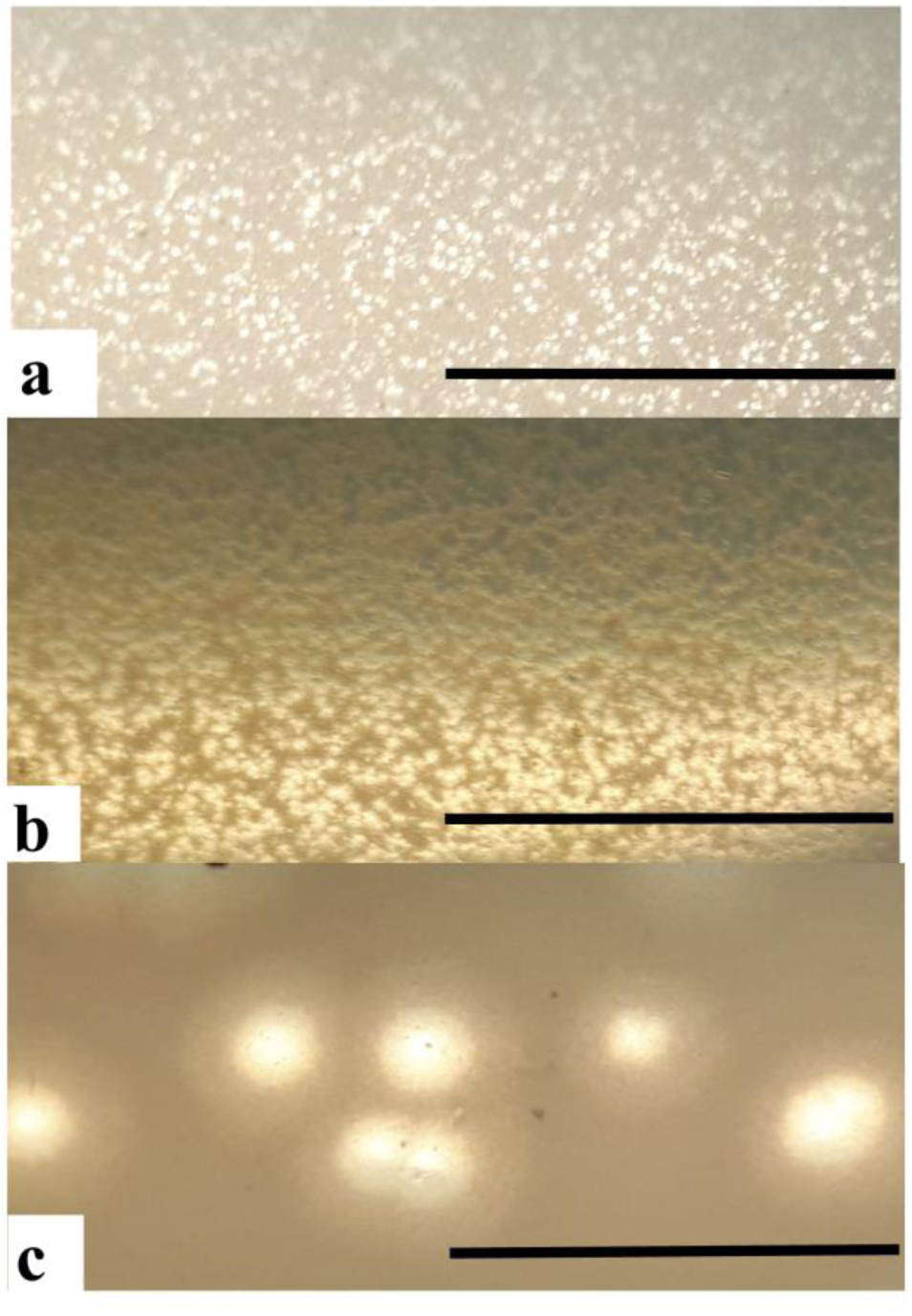
Morphology of phage plaques from the primary isolate (a) and their ability to cause a confluent lysis of the bacterial lawn (b) comparing with morphology of phage T4wt plaques (c). The experiment was conducted in serial repetitions on a lawn of the parental host *E. coli* strain. The bar equals to 1 cm at each photo.

Further EM examination showed that the primary phage isolate contained T4-like particles having contractile tail and elongated icosahedral head, thus belonging to *Myoviridae* family and A2-morphotype, respectively (hereinafter ‘myovirions’). Thus, based on the features of the phage negative colonies it was assumed that the studied phage isolate contained an attenuated viral population.

### 3.2 The chromatographic profile of the primary phage isolate

A chromatographic profile of UV-adsorbing contents of the primary phage isolate consisted of three main peaks eluted by 0.15, 0.25, and 0.4 M NaCl step gradient (Fig 2, a). In the case of peaks II and III, the presence of several poorly resolved peaks can be expected according to the curves A260 (black), A280 (red) and the fluorescence intensity curves (green). Based on the electrophoretic separation of the native peak fractions (Fig 2, b, c), plaque assay, and EM examination we found that the start of phage elution corresponded to zones of local minimums on the curve A260/280 (blue), and phage enriched peaks can also be identified by this ratio.

**FIGURE 2.**
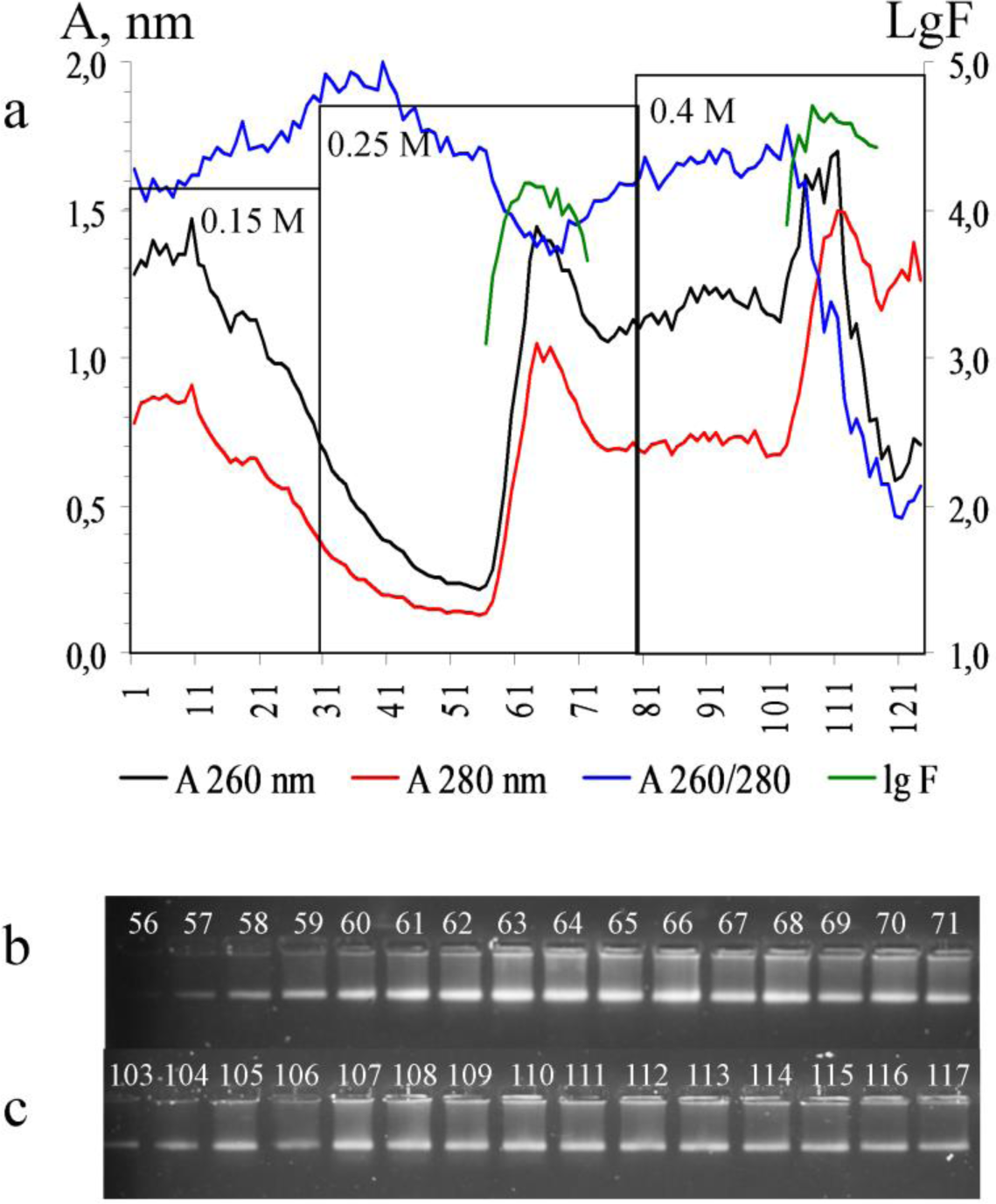
Chromatographic profile of the primary phage isolate (a) and detection of native phage capsids in agarose (b, c). The fractions were eluted by a stepwise gradient of 0.15, 0.25, and 0.4 M NaCl. The peak maximums coincided both at 260 and 280 nm. a - 0.15, 0.25, and 0.4 MNaCl – the elution steps; A, nm – UV-absorbance, lgF – fluorescence intensity logarithm. b, c - separation of native fractions of peaks II and III, respectively, in agarose gels. Capsid mobility was similar in samples of peaks II and III (peak I fractions were not analyzed). Fractions 63, 64 (0.25 M NaCl), and fractions 105 and 107 (0.4 M NaCl) had the most intensively fluorescent bands.

Electron microscopy examination has shown that all the three main peaks contained myovirions (Fig 3, a). A part of them had specific defects in the tail sheath that may additionally indicate that the studied myophage pool was mono-species. Surprisingly, peak III contained also a substantial amount of virions belonging to the *Siphoviridae* family (siphovirions, Fig 3, b). However, aside from the above-described pinprick-type colonies, no morphologically distinct plaques indicating of a mixed phage sample were observed.

**FIGURE 3.**
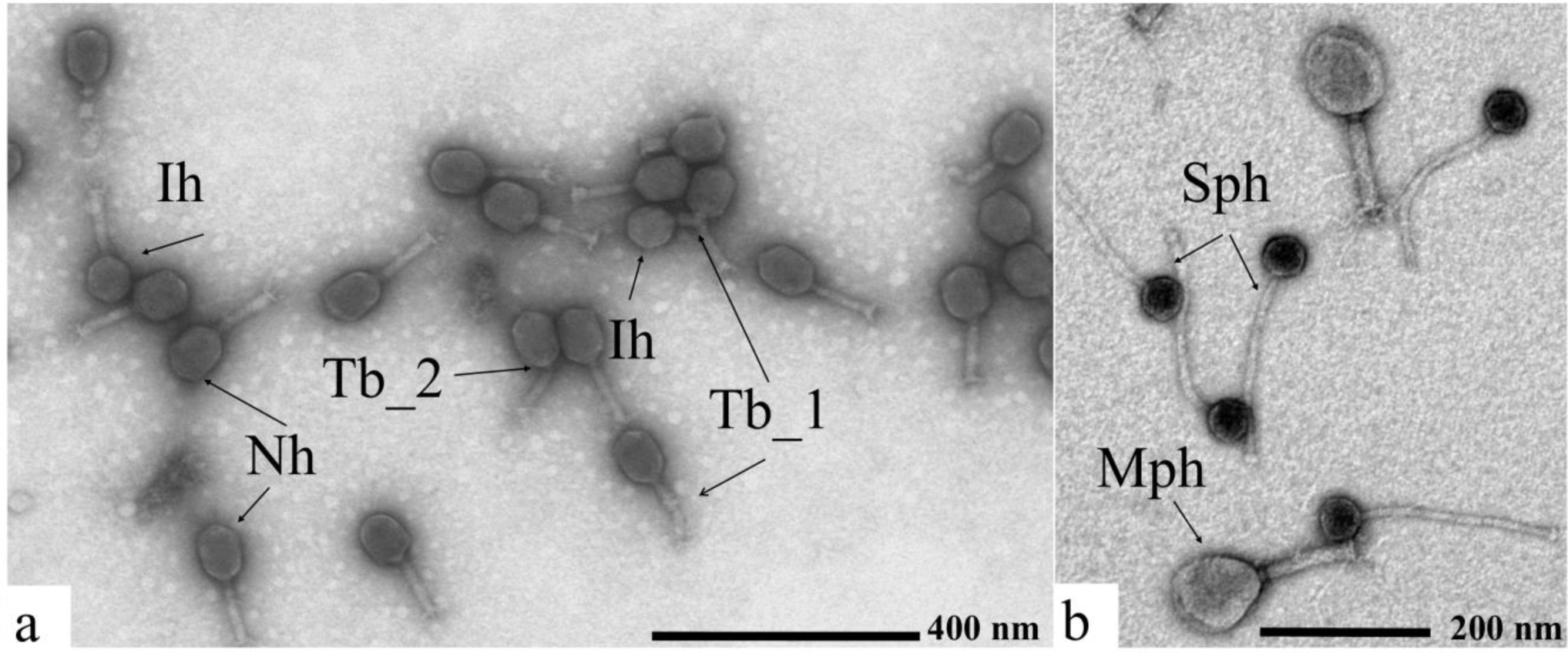
Gross morphology of phage particles eluted in peak I (a) and peak III (b). Peak I consisted of myovirions only, while peak III contained siphovirions additionally. Peak II contained morphologically homogenous myovirions that were similar to other samples (data not shown). The bars are equal to 200 nm. a – “Nh” – normal elongated T4-like head; “Ih” – isometric head, a naturally occurring aberrant form of phage T4 capsid; “Tb_1” and “Tb_2” – specific tail breaks observed within the myovirions. b – “Sph” – *Siphoviridae* virions, “Mph” – *Myoviridae* virions

### 3.3 Restriction patterns of the chromatographic peaks

Despite the fact, that peak III fractions contained a fairly large number of siphovirions with filled heads (appeared as dark capsids on EM-micrograph, Fig 3, b), a comparative restriction analysis of total phage DNA revealed no difference in the corresponding patterns (Fig 4). The patterns appeared to be identical to those of phage Lw1, which had been isolated from this primary phage pool earlier [3]. To disprove a speculation that virion nucleic acid of the siphophages was destroyed during manipulations, TAFE-separation of total phage DNA from the same samples was conducted.

**FIGURE 4.**
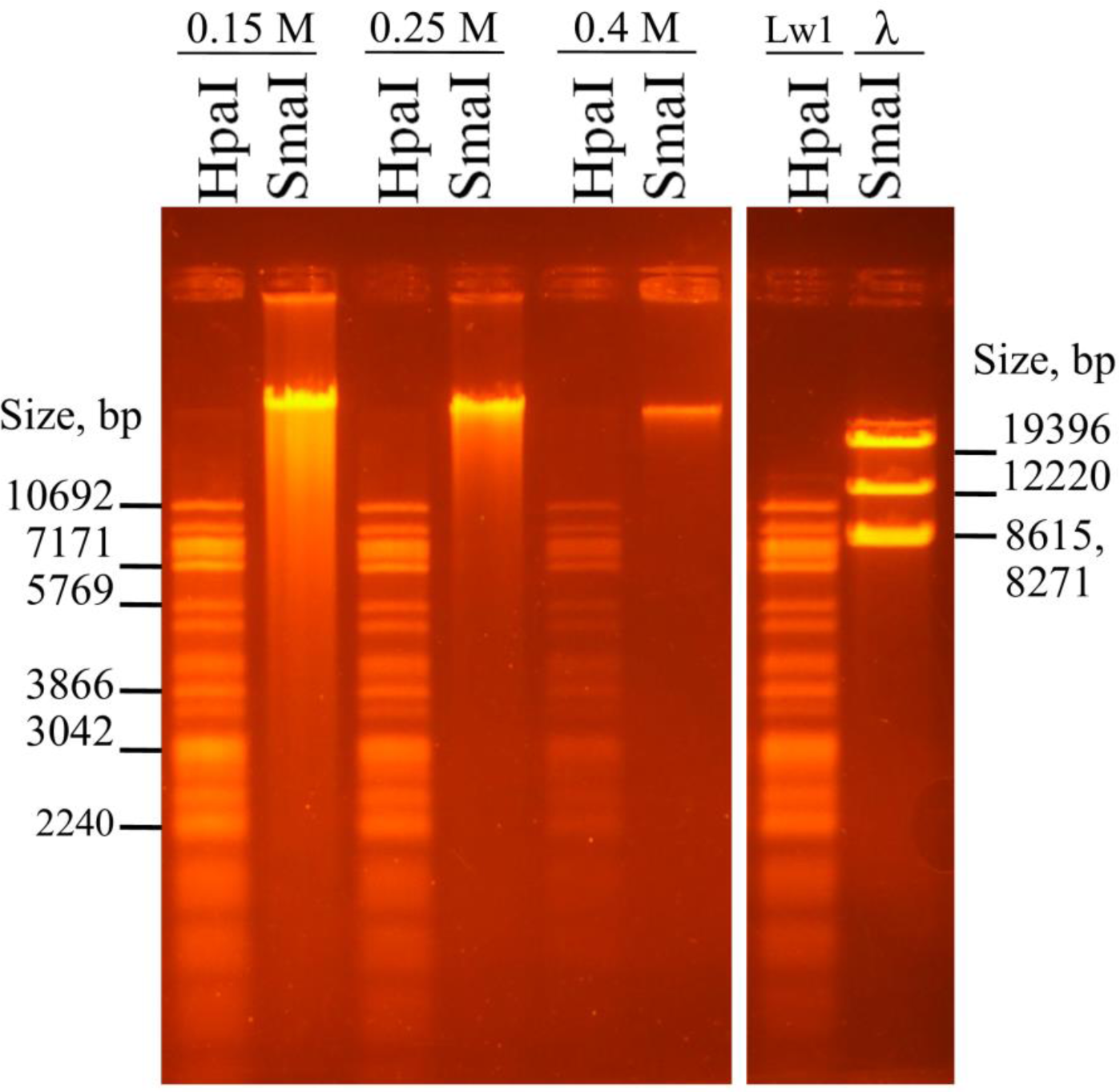
Restriction analysis of total phage DNA. isolated from chromatographic peaks I (0.15 M NaCl) and II (0.25 M NaCl), which contained myovirions only, and DNA from peak III (0.4 M NaCl) containing a mix of myo- and siphovirions in comparison with restriction pattern of Lw1 phage DNA.

### 3.4 Transverse altering field gel electrophoresis (TAFE) of total phage DNA from the chromatographic peaks

TAFE-analysis (Fig 5) showed that the primary phage isolate consisted of virion DNAs of three size classes. The DNA-genomes of approximately 170 (I class) and 120 (II class) kb were found in samples of all chromatographic peaks. Evidently, they corresponded to the observed elongated icosahedral heads and rare isometric heads of the primary Lw1 virions (Fig 3, a).

**FIGURE 5.**
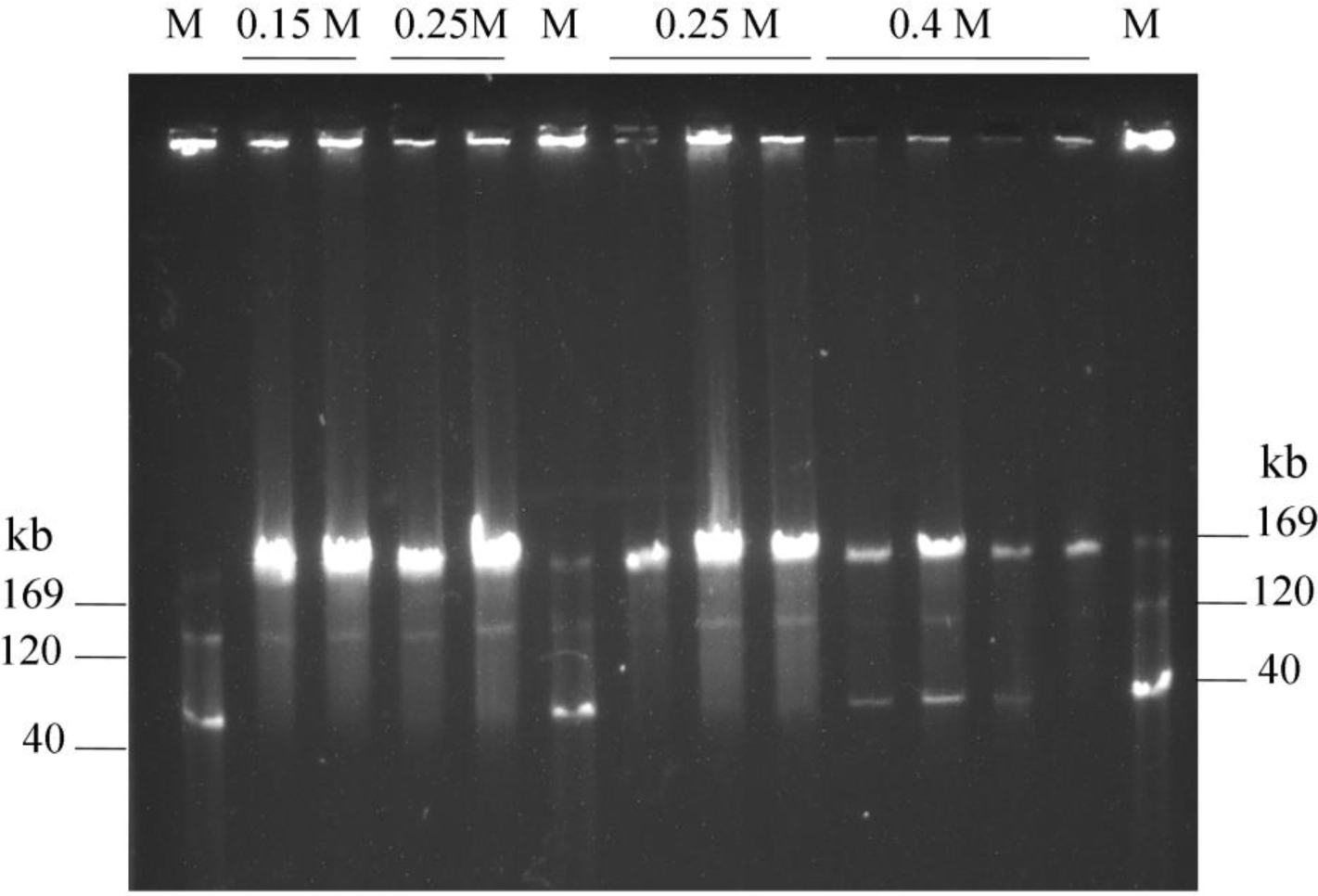
TAFE of total DNA samples extracted from chromatographic peaks I, II, and III. Samples were tested in different concentrations. M – marker composed of 169 kb virion DNA of phage T4, 120 kb virion DNA of phage T5, and 40kb virion DNA of phage T7, isolated by the same technique as described for the TAFE-phage samples.

It is well known that isometric heads are common in phage T4 populations [12]. Their virion DNA size is about 2/3 of a complete phage T4 genome [13] that is approximately 113 kb. In our case, 120 kb was about 2/3 of a complete genome length of phage Lw1 that is 176 227 bp [3]. Structural similarities of basal plates, as well as a specific defect in some of phage tail sheaths (data not shown), allowed to identify them as analogous petite-phages of the prevailing myophage population.

Mixed phage sample of the peak III contained additional DNA molecules of 40 kb length (III class). As anticipated, this 40 kb DNA was not found in the samples of peaks I and II. Therefore, the siphovirions release nucleic acid of size correlating with their head diameter (see below).

### 3.5 Morphological analysis of the myophage population

To identify differences in the myovirions structure that correlate with their chromatographic behavior and to reveal the population morphological diversity we analyzed phage tail length, capsid height and width. The three chromatographic peaks I (0.15 M), II (0.25 M), and III (0.4 M) are considered hereinafter as three subpopulations I, II, and III of the primary myophage Lw1, respectively.

Gross morphology of myophages eluted in different peaks is presented in Fig 6, a–c. The low values of dispersions (Table 1) suggested that myovirions from the main subpopulation II were more uniform by their size than those ones from minor subpopulations I and III.

**Table 1.** Indexes of dispersion* of the measured linear parameters of the *Myoviridae* virions from the chromatographic subpopulations

| # | Subpopulation | I (0.15 M) | II (0.25 M) | III (0.4 M) |
| --- | --- | --- | --- | --- |
|  | Linear parameter of virion |  |  |  |
| 1 | Tail length | 21.7 | 7.2 | 12.7 |
| 2 | Tail sheath length | 27.7 | 11 | 14.5 |
| 3 | Basal plate front height | 9.7 | 5.1 | 7.9 |
| 4 | Capsid height | 37.5 | 20.7 | 24.9 |
| 5 | Capsid width | 27.3 | 23.8 | 18.4 |
| 6 | Coefficient of capsid elongation | 0.01 | 0.006 | 0.005 |

\*data number for each parameter was >150; standard error of the mean, nm – 0.23 – 0.4.

**FIGURE 6.**
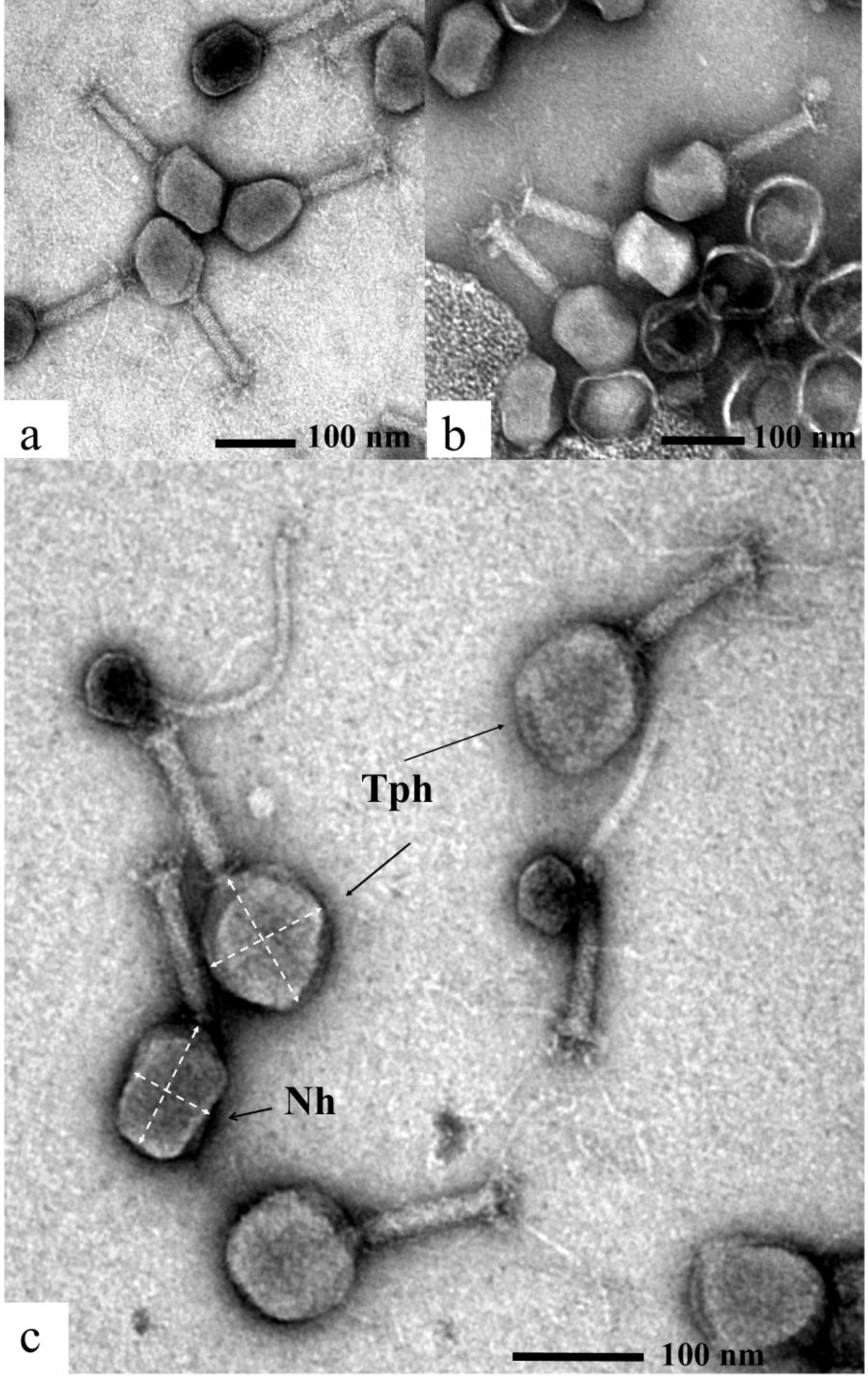
Gross morphology of particles from subpopulations of the primary phage Lw1 isolate. a, b, and c – morphology of myovirions from peaks I, II, and III, respectively.”Nh” – normal elongated icosahedral capsid; “Tph” – tubby-phages, the myovirions featuring a capsid shape close to spherical in comparison with the elongated icosahedral head.

Modal dimensions of the myovirions eluted into different chromatographic peaks are presented in Table 2. The complete tail length was 124 nm for the major part of particles (Supporting fig 1). There were some differences between the subpopulations in the tail sheath length and the frontal height of the basal plate that, generally, did not affect the complete length of the tail. These differences can be explained by discrete conformational alterations of the tail structural proteins due to different concentrations of NaCl in the samples. Thus, the tail size had no impact on morphological diversity of the myophage population and we believe that it could not significantly affect the affinity of the virions to DEAE-cellulose.

**Table 2.** Titres and morphological parameters of primary population of phage Lw1.

| Chromatographic peak<br>(subpopulation) | Phage titer,<br>PFU/ml | Complete tail, nm | Tail sheath, nm | Basal plate, nm | Capsid, height x width, nm | Aspect ratio* | Capsid surface area, nm<sup>2</sup> | Capsid volume, nm<sup>3</sup> |
| --- | --- | --- | --- | --- | --- | --- | --- | --- |
| I, 0.15 M NaCl | 10<sup>9</sup> | 124 | 111<br>114 | 13 | 100x72 | 1.36 | 21.4x10<sup>3</sup> | 29.4x10<sup>4</sup> |
| II, 0.25 M NaCl | 10<sup>12</sup> | 124 | 109 | 13 | 111x78 | 1.42 | 23.2x10<sup>3</sup> | 31.8x10<sup>4</sup><br>34.9x10<sup>4</sup> |
| III, 0.4 M NaCl | 10<sup>10</sup> | 123 | 109 | 12<br>15 | 102x81<br>102x84 | 1.16<br>1.20<br>1.24 | 23.9x10<sup>3</sup><br>26.0x10<sup>3</sup> | 35.1x10<sup>4</sup><br>38.2x10<sup>4</sup> |

\* - aspect ratio is capsid height/capsid width

However, the three myophage subpopulations differed in capsid dimensions and their sizes correlated with the eluant ion strength. The modal capsid size of subpopulation II was 111×78 nm (Table 2, Supporting fig 2). This capsid dimensions reflect the whole primary Lw1 population, since the subpopulation II contained the highest phage titer, 10^12^ PFU/ml. Virions of subpopulations I and II differed in their capsid modal sizes, but both had elongated icosahedral heads with the modal aspect ratios (capsid height/width) of 1.36 and 1.42, respectively.

For capsids of subpopulation III three discrete modal aspect ratios were found (Table 2, Supporting fig 2). However, these values were close to 1 and lower than those ones of elongated heads that indicated these capsids were rather isometric than elongated. Fig 6, c shows that the virions gathered in the peak III (0.4 M) indeed had capsids with a sphere-like shape clearly distinguishing from the elongated icosahedral head. These aberrant particles had restriction patterns identical to that of phage Lw1 (Fig 4) and were named tubby-phages because of the specific shape of their capsids. Ranged data series additionally demonstrated the differences in a sample of individually calculated values with no data averaging and whether the dimensions of each subpopulation could be collected in separate plurality. Absence of a significant difference in the phage tail length between the phage subpopulations is demonstrated by the corresponding ranged data series on Fig 7, a. Nevertheless, the ranks of the capsid height, width, and aspect ratios show convincing distinction and confirm that heads of myophages of the three subpopulations differ sufficiently (Fig 7, b-d). Thus, it is logical to assume that surface area and volume of myophage capsids should differ too.

**FIGURE7.**
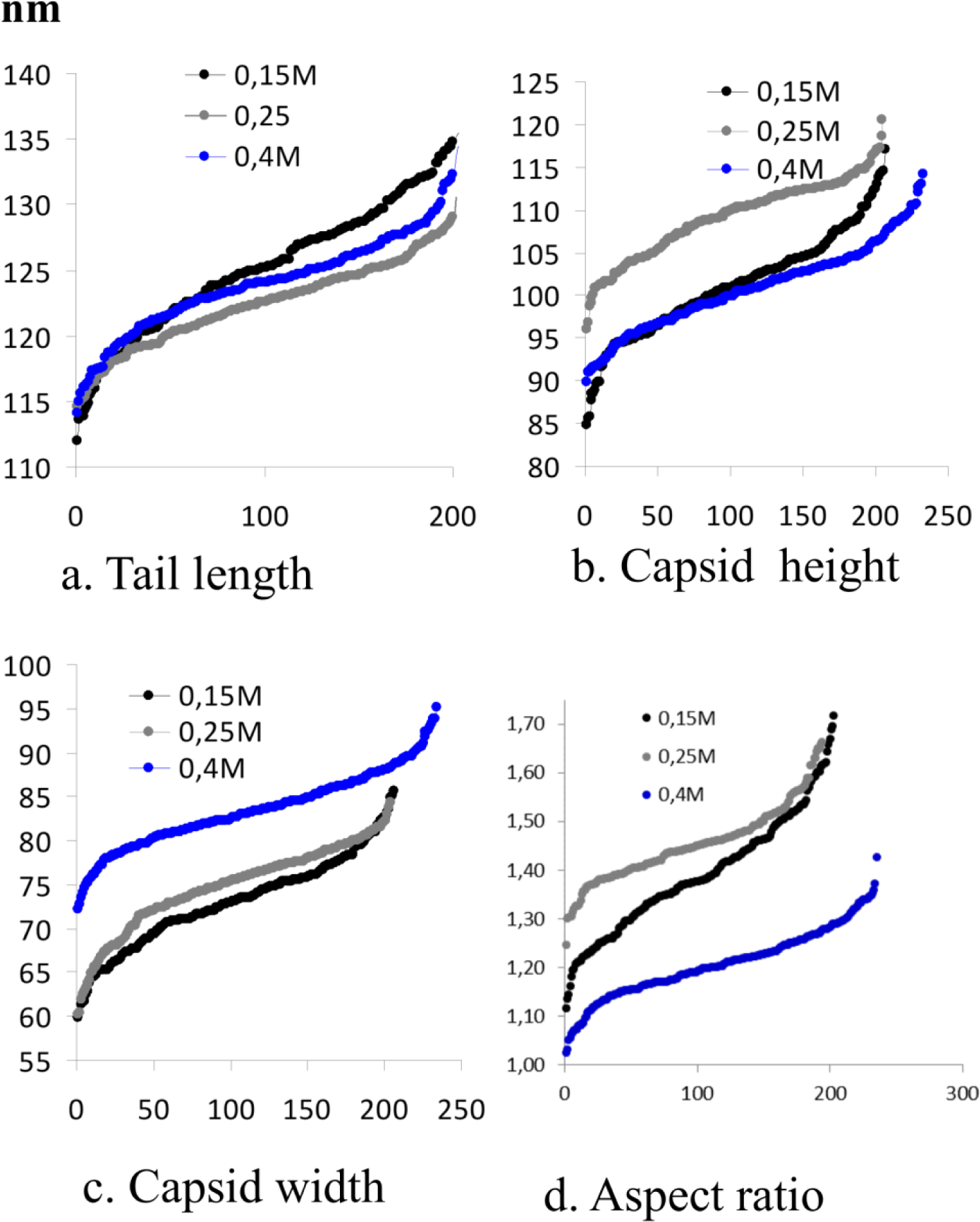
Ranged data series for *Myoviridae* virions from the main chromatographic peaks. n – data number.

#### 3.5.1 Conditional capsid surface area and volume of the myophage capsids

Ranged data series of conditional myocapsid surface areas S and conditional volumes V are presented in Fig 8. They provide a clear confirmation of the existence of the three myovirion subpopulations that differed by their capsid surface areas and volumes. As shown in the diagrams, the difference in the values of the discussed parameters were sufficient, therefore data of each subpopulation aligned into discrete sequences.

The modal values of S and V for the myophage capsids are presented in Table 2 (Supporting fig 3 and 4). Accepting these values for subpopulation II being equal to “1” (since it contained virions with the representative for the whole myophage population capsids), S and V values were calculated to differ by around ±10-20% in subpopulations I and III.

**FIGURE 8.**
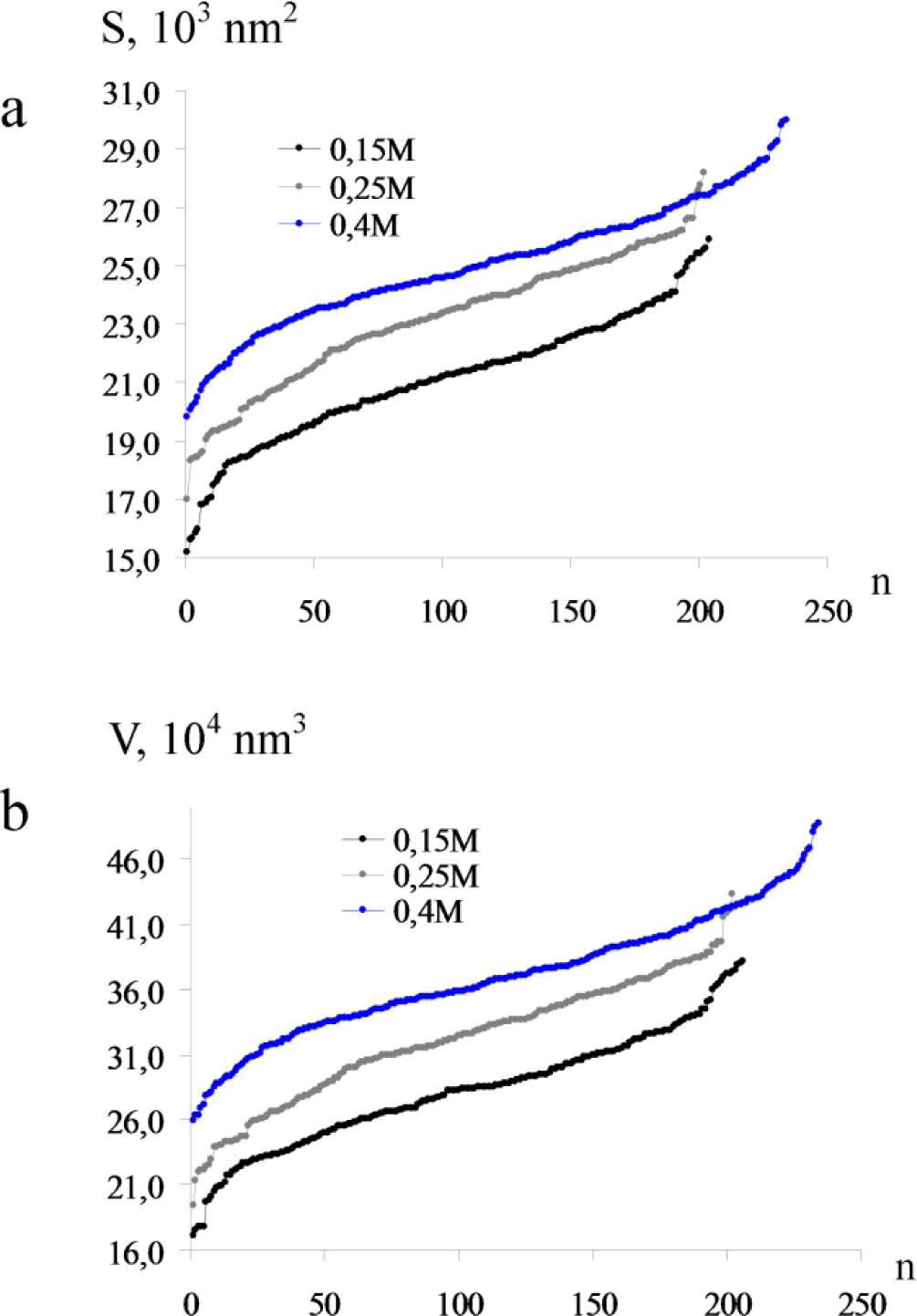
Ranged data series of conditional capsid surface area S (a) and conditional capsid volume V (b) of the myovirions. n – data number.

### 3.6 Morphological analysis of siphovirions

During EM examination, two types of *Siphoviridae* virions of B1 morphotype (particles with non-contractile tails and isometric heads) were observed. The prevailing part of them (Fig 9, a) had a modal capsid diameter of 51 nm (Fig 9, d) and the modal phage tail length of 215 nm (Fig 9, e). In some cases, it was possible to scrutinize the tail attachment apparatus structure of these phages (Fig 9, b). The second type of siphovirions was represented by rare particles with over-long tails of 300-400 nm length attached to empty capsids (Fig 9, c). In this case, the capsid might play the main role in negative virion surface charge of siphophage population, while the phage tail length suggests its intrapopulation heterogeneity or the presence of two phage species.

**FIGURE9.**
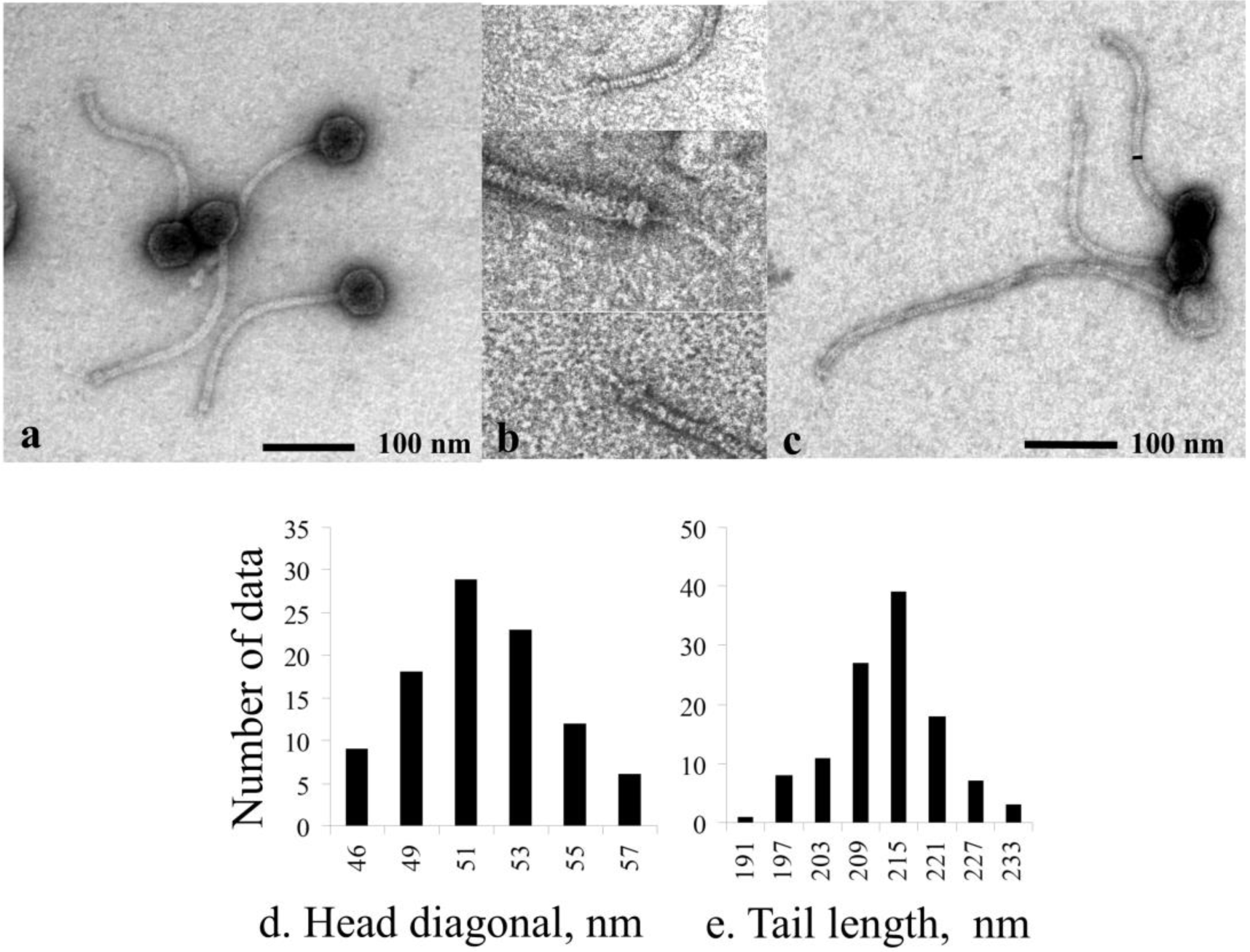
Gross morphology and frequency distribution diagrams of *Siphoviridae*virions from the primary isolate. a – commonly observed siphovirions with tails of 215 nm length. b – basal plate of siphophages with 215 nm tails. It consisted of two-disc basal plate and single central fibril. The collage was combined from fragments of micrographs captured with different magnification. c – rare siphovirions with extremely long tails of 300–400 nm length. d – a frequency distribution diagram of the *Siphoviridae* phage capsid diameter. Only one modal capsid diameter equal to 51 nm was found. e – a frequency distribution diagram of the *Siphoviridae* phage tail lengths. One modal length of 215 nm for non-contractile tails was identified statistically. The extra-long tails were not included to the reckoning.

## 4 Discussion

Intrapopulation morphological diversity, or heterogeneity, of phage T4, that is a prototype of the whole *Myoviridae* phage family in order *Caudovirales*, is well known. The virion of phage T4 is a model of tailed viruses of A2-morphotype [14]. The normal capsid of phage T4 is an elongated icosahedron and its molecular structure recently was studied in details [15]. Five types of clearly distinguishing aberrant (abnormally shaped) capsids with packed DNA are known for T4 phage. There are three types of capsids with alternating length, but the same width such as: isometric (small icosahedral capsids containing about 2/3 of T4 genome) [12, 13, 16], intermediate (shorter by 4-5 nm, but the geometry of the capsid resembles normal elongated icosahedron) [17, 18], and giant T4 heads (elongated by 1.5 times or more comparing to normal T4 capsid) [19]. Two capsid aberrations affecting its width are known such as biprolate heads – protruded on one side of a fivefold axis, and triprolate capsids – protruded on both lateral sides [20].

The aberrant capsids assembling during the T4wt infection are supposed to be rare [21], but their formation can be induced by amino acids and anti-metabolites [22], or be a result of specific mutations (amber-mutants) [23]. Naturally occurring capsid aberrants in T4wt phage are known only as those ones having isometric and biprolate geometry. The isometric T4 virions are non-infectious, since they contain only about 2/3 of the whole phage genome [13] and their increased amount in the lysate correlates with the lower recombination frequency of T4 mutants [24]. The naturally occurring biprolate phages are more frequently assembled with two or more tails, than with one [12]. As the giant phages, they could be infectious since they contain DNA of more than one T4 genome length [25], but it is unknown how the additional tails function during adsorption and DNA injection [20]. By analysis of amber and *ts*-mutations it was shown that twelve genes are involved in T4 capsid morphogenesis [26]. However, only two of them, genes *22* and *68* [20, 27] are involved in control of capsid width.

The mechanisms of natural occurrence of the aberrant capsids are unknown, though, it was long ago observed, that, in T4 and T2 phages, the ratio of isometric, multi-tailed and normal virions depends on the multiplicity of infection and/or physiological state of the culture [24, 28, 29].

The observed tubby-phages in phage Lw1 primary population were symmetrically protruded. They also can be referred to triprolate aberrant virions, but this needs to be proven by additional studies. Besides, they were not observed to be assembled with multiple tails, while about one thousand virions had been examined in general. The primary Lw1 virions gathered in the peak I can be considered as having intermediate capsid that retains the geometry of normal head. In our case, all the three primary phage Lw1 subpopulations were infectious, thus it is interesting how the capsid size affects on the virion DNA structure, and phage-to-phage or phage-host genetic exchanges. Also, it is unclear, whether the plaques of the subpopulation I do reflect the development of myovirions with intermediate capsids mostly gathered to this peak, or they arise due to propagation of the ‘normal’ Lw1 virions randomly eluted into the peak I fractions.

Using phage T4 as benchmark against which T4-like phages isolated from environment were analyzed, many viruses producing aberrant capsids and tails in laboratory conditions were reported [30, 31]. Such aberrations are considered as results of natural mutations in genes homologous to the corresponding T4 sequences. Thus having an attenuated Lw1 population should be an ordinary situation, rather than isolation of “ideal” lytic phage.

As it was noted by Rao&Black [32] even for well-studied phage capsids the problem of capsid volume determination was elaborated insufficiently. Moving towards understanding of mechanisms of phage Lw1 intra-population diversification we conducted a series of preliminary experiments on the impact of host cell on alterations in the phage population structure. Stocks of Lw1 and T4 obtained on *E. coli* B^E^ were used to infect cells of *E. coli* B^E^, BL21 and BL21(DE3). The obtained lysates were subjected to chromatography by the protocol described in Materials and Methods. In Fig 10 it is shown that height and localization of peaks on chromatographic profiles of phage T4 from different hosts were almost the same, but the profiles of phage Lw1 had severe differences.

**FIGURE 10.**
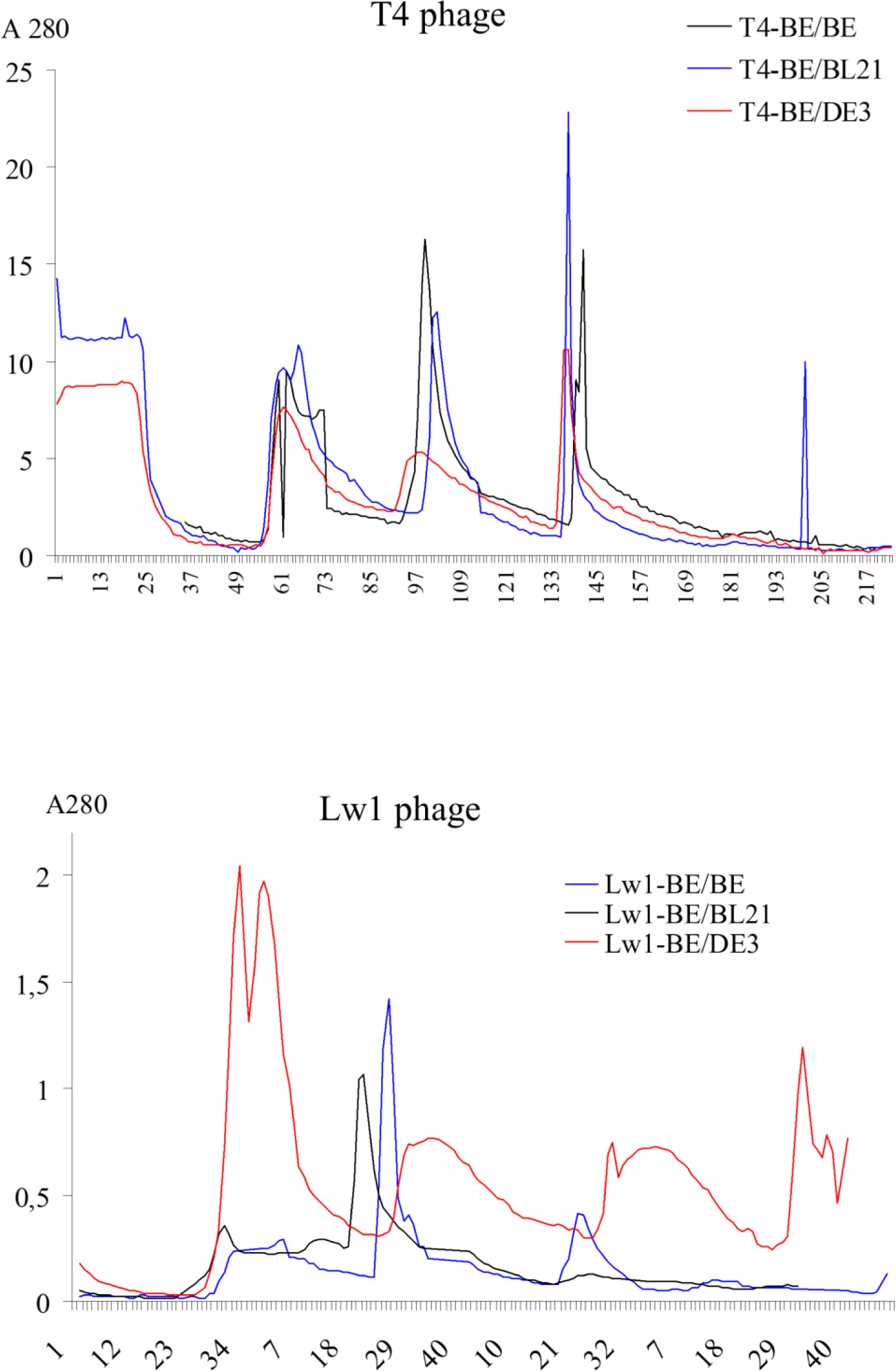
Chromatographic profiles of phages T4 and Lw1 populations obtained on *E. coli* B^E^ (black), BL21(blue) and BL21(DE3) (red).

The profile of Lw1 propagated on BL21(DE3) was close to that one obtained for its primary population and described in this paper, while Lw1 profiles from the other host strains contained only one main peak gathered by 0.25 M NaCl. Despite the fact, that the peak number stays, generally, the same, the volumes of the virion-containing fractions of phage Lw1 change significantly depending on the host bacterium. Thus, it is more probable that intrapopulation diversity of phage Lw1 is related to both phage and host genetics, because the structure of phage T4 population stays the same on different hosts.

Each case of naturally occurring phage lysis should demonstrate a specific pattern comprising a fixed proportion of prevailing and aberrant virion forms, as well as non-assembled virion components. This ratio usually suggests about intensity and stage of viral infection, as well as about host permissivity to a particular virus. In this context, the question of phage Lw1 attenuation remains open. It is unclear, what led to the virions heterogeneity and attenuation – contamination of the batch-culture by already defective virus or a spontaneous mutation that happened in the bio-reactor. Also, it is unclear whether the observed heterogeneity was related to the attenuation and what was a role of original host and/or the phage itself in developing of the attenuation.

Detection of multiple morphological and genetic forms of viral particles in population heterogeneity studies is complicated due to the absence of precise universal screening system for a large number of virions, some of which may have lost their infectivity due to the reproduction peculiarities or conceal it under proposed experimental scenarios. In these cases, AEC may serve as an effective tool for scrutinizing the phage population structure by the density of virion net surface charge since both these features can be regarded as universal and effective selection markers suggesting the occurrence of different changes in phage genetics and virus-host interactions. The main goal of the proposed research was achieved by AEC on fibrous DEAE-cellulose as the principal method allowing for the analysis of morphological (structural) heterogeneity of naturally occurring phage populations before the isolation of pure phage lines at the first step of the research. Today, there are commercial chromatographic columns available that allow for purification and separation of phage lysates, but they have quite high prices and maximal reported volume of phage suspension concentrated by these tools was 470 ml [33]. Since we needed to treat a sample of 5 liters volume avoiding high-cost tools and logistic challenges, the classical approach to anion-exchange chromatography was applied. This approach is routinely used in our lab and its high efficacy had been demonstrated [34]. As it was shown above, the used anion exchanger allowed us to not only purify and concentrate a big volume of the phage sample, but effectively isolate and enrich the hidden population of the siphophage as well as the minor sub-populations of the myophage.

As this study has demonstrated, lysis of an industrial recombinant *E. coli* strain may be related to several bacteriophages at the same time. We tend to think, that the siphophage gathered in one peak was a single species. The observed over-long tails rather were aberrant structures, since only a few such virions were found among a number of others. Such tail aberrations are known for *Siphoviridae* viruses. For example, *Salmonella* phage MB78 produced 3-4% of over-long aberrant tails [35].

Biodiversity of viral populations is under close attention since viruses are the most numerous living entities accompanying cellular organisms. Despite the extreme undesirability, sporadic phage lysis at biotechnological facilities could be regarded as a quite convenient semi-natural system, or even model, to study phage population ecology. In the described case of phage lysis, a competitive exclusion between the populations of sipho- and myophage evidently existed. This fact is interesting because based on plaque morphology and ability to propagate in subsequent passages we concluded that the prevailing myophage population was attenuated. By the methods used we did not identify any common specific defect in the myovirions structure that might explain such a phage behavior. However, it was found that this defect, which did not allow the general phage pool to produce robust negative colonies on solid LB-medium, was overcome by an increased multiplicity of infection of the sensitive cells. In such context, sporadic phage lysis of *E. coli* BL21(DE3) and its recombinant derivatives in large-scale bio-reactor is one of the most suitable models. Advantages of this host-virus system are massive knowledge of *E. coli* physiology, genetics and genomics, as well as of its lytic and temperate phages. The investigation of cases of industrial lysis cases may lead not only to the developing of ways for prevention of phage contamination, but also to the evaluation of the complexity of semi-natural phage populations and understanding of the key mechanisms, which affect the development of viral genetic and morphological diversity, virus-like particles forming processes, attenuation of virus population and ecological roles of bacteriophages in natural microbial communities under different types of phage infections.

## Acknowledgment

The authors appreciate N. Kozyrovska, L. Gorb, M. van Raaij, M. Zlatohurska, I. Faidiuk, and O. Glushko for discussion and help with the manuscript preparation. The research was funded by the National Academy of Sciences of Ukraine (http://www.nas.gov.ua, Grant number 0115U004128).

---

**FIGURE S1.**
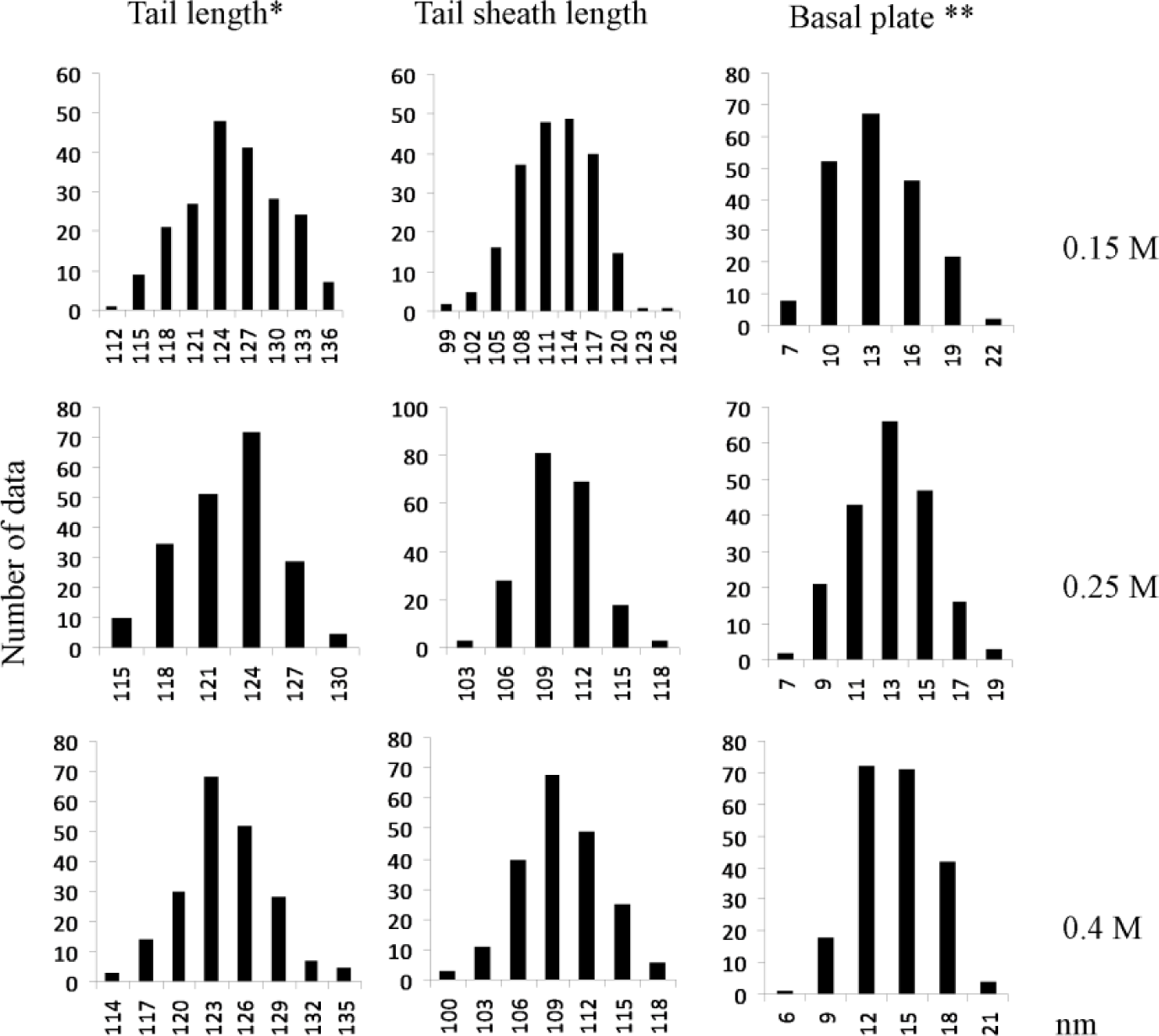
Frequency distribution diagrams of *Myoviridae* phage tail linear dimensions according to their chromatographic subpopulations. * “tail length” means length of a complete phage tail from the start of phage neck till the visible distal end of the basal plate; ** frontal height of the basal plate was calculated as difference between the complete phage tail length and tail sheath length

**FIGURE S2.**
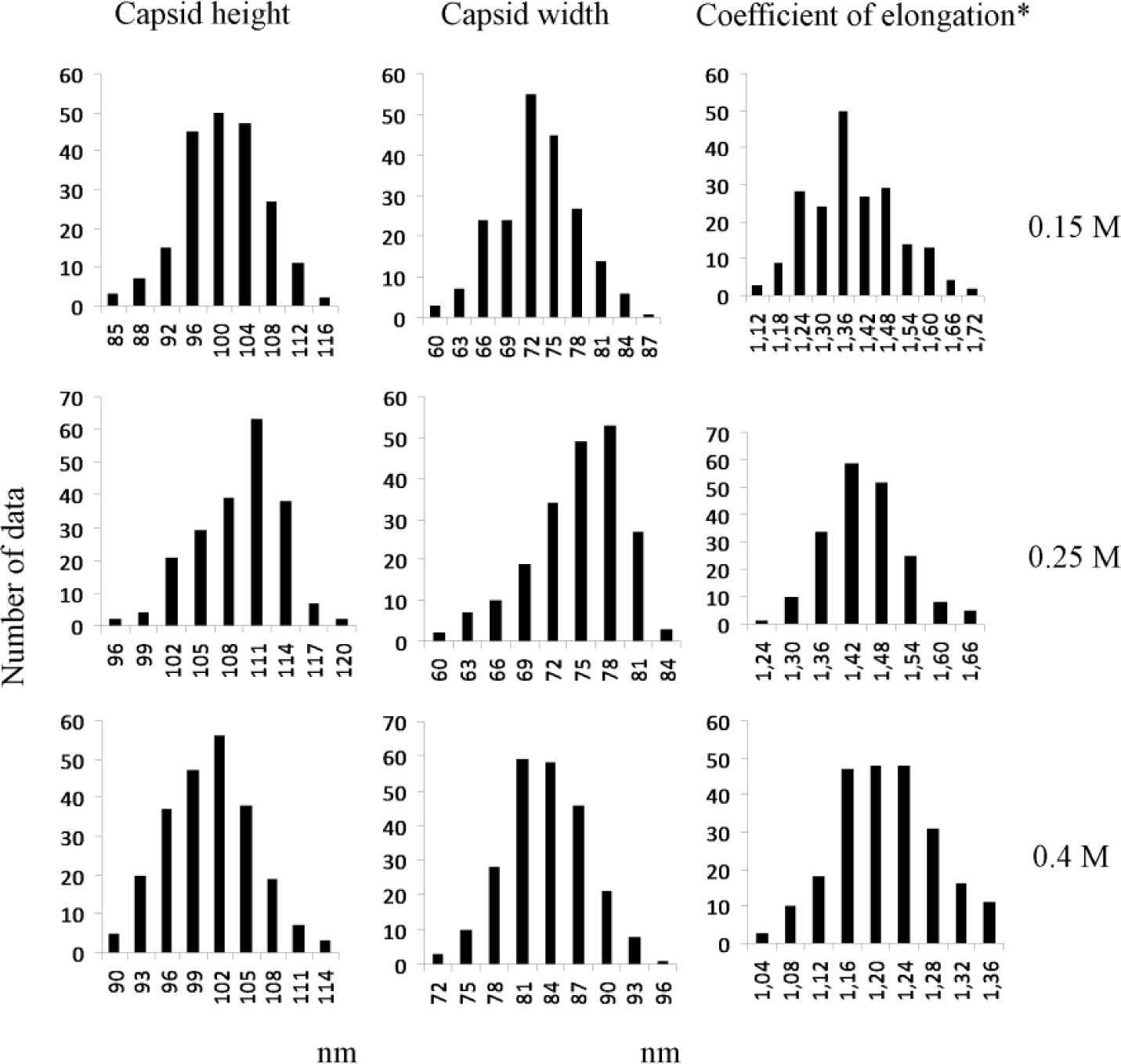
Frequency distribution diagrams of *Myoviridae* phage capsid linear dimentions according to their chromatographic subpopulations. * coefficient of the capsid elongation was calculated as ratio of the capsid height to the capsid width

**Fig S3.**
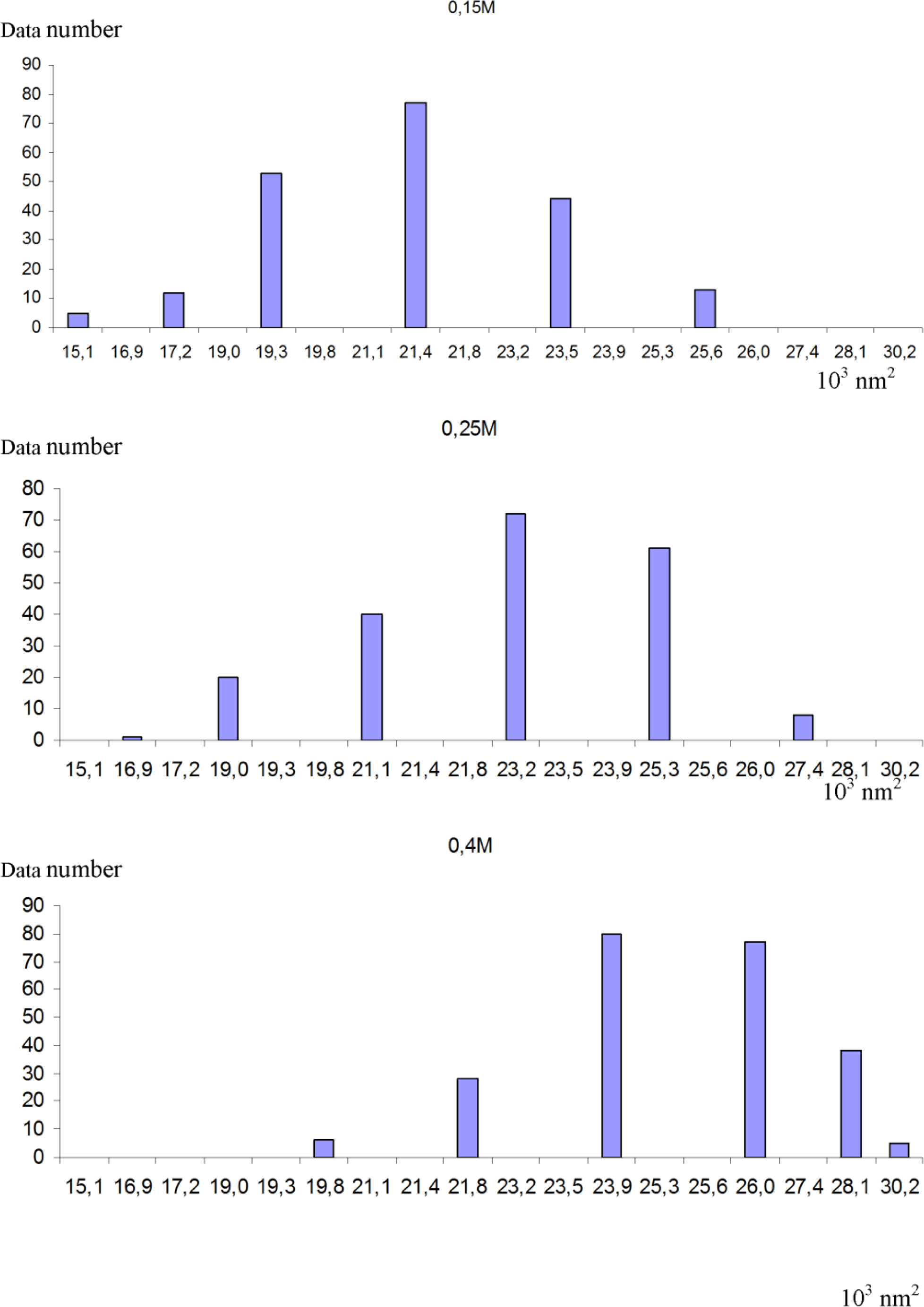
Frequency distribution diagrams of conditional capsid surface square S.

**Fig S4.**
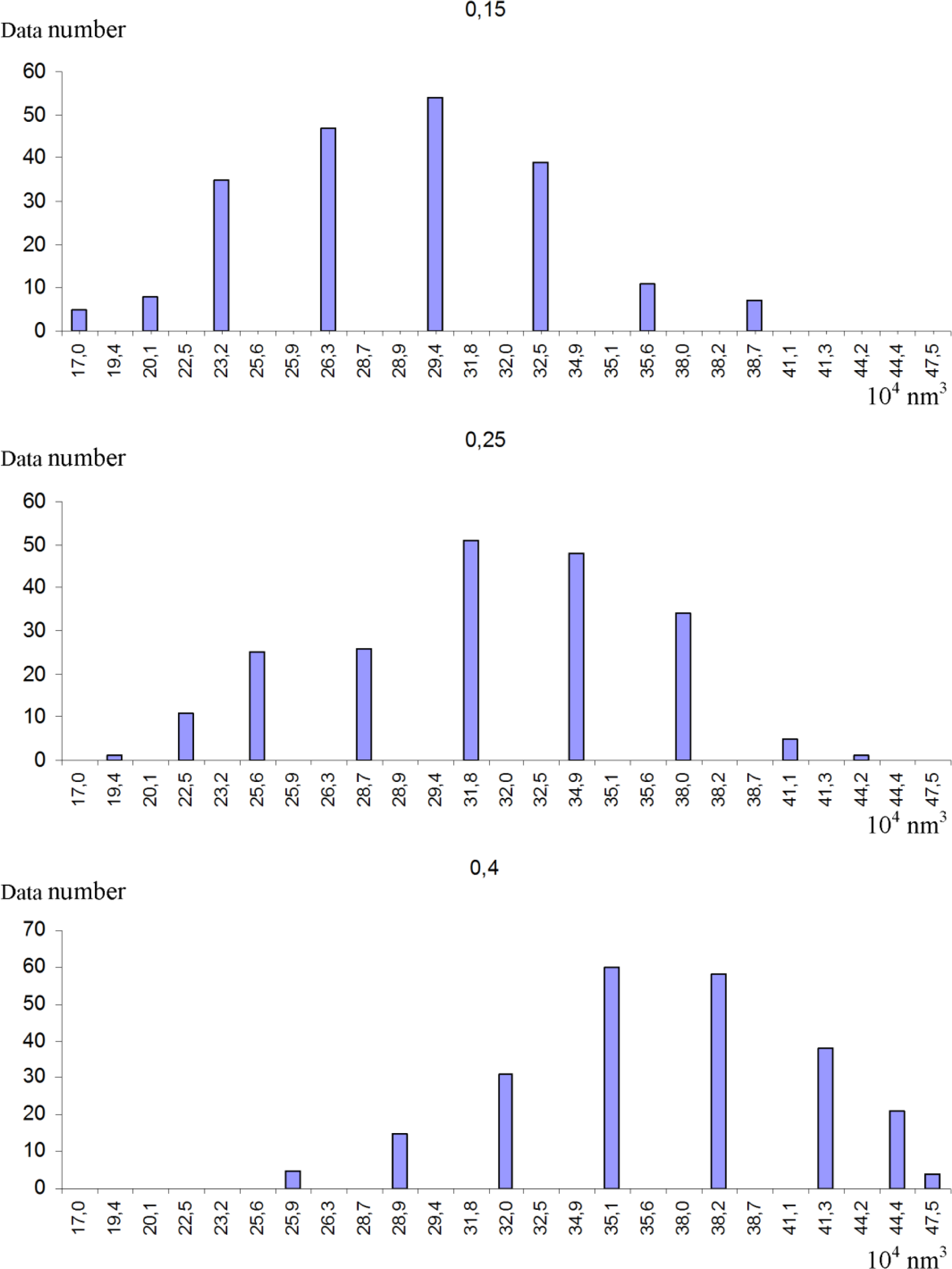
Frequency distribution diagrams of conditional capsid volume V.

## References

[1] Studier FW, Moffatt BA. Use of bacteriophage T7 RNA polymerase to direct selective high-level expression of cloned genes. J Mol Biol 1986;189:113–30.

[2] Li S, Liu L, Zhu J, Zou L, Li M, Cong Y, et al. Characterization and genome sequencing of a novel coliphage isolated from engineered *Escherichia coli*. Intervirology. 2010;53:211–20.

[3] Kushkina, AI, Tovkach FI, Comeau AM, Kostetskii IE, Lisovski I, Ostapchuk AM, et al. Complete genome sequence of *Escherichia* phage Lw1, a new member of the RB43 group of pseudo T-Even bacteriophages. Genome Announc. 2013;1(6), e00743–13.

[4] Kropinski AM, Mazzocco A, Waddell TE, Lingohr E, Johnson RP. Enumeration of bacteriophages by bouble agar overlay plaque assay. In: Clokie MR, Kropinski AM, Editors. Methods in molecular biology, vol 501. Bacteriophages. Humana Press; 2009.

[5] Tikhonenko TI, Koudelka IA, Borishpolets ZI. Phage concentration and purification by the column chromatography method. Mikrobiologiia. 1963;32:723–6.

[6] Sambrook J, Fritsch EF, Maniatis T. Molecular cloning: a laboratory manual. 2^nd^ ed. Cold Spring Harbor Laboratory Press; 1989.

[7] Serwer P, Pichler ME. Electrophoresis of bacteriophage T7 and T7 capsids in agarose gels. J Virol. 1978;28:917–28.

[8] Serwer P, Hayes SJ, Thomas JA, Griess GA, Hardies SC. Rapid determination of genomic DNA length for new bacteriophages. Electrophoresis. 2007;28:1896–902.

[9] Schneider CA, Rasband WS, Eliceiri KW. NIH Image to ImageJ: 25 years of image analysis. Nat Meth. 2012;9:671–5.

[10] Sturges, HA. The choice of a class interval. J Am Stat Assoc, 1926;21:65–6.

[11] Zwillinger D. CRC standard mathematical tables and formulae. Chapman and Hall/CRC, 2002. p.220.

[12] Boy de la Tour E, Kellenberger E. Aberrant forms of the T-even phage head. Virology. 1965;27:222–5.

[13] Eiserling FA, Geiduschek EP, Epstein RH, Metter EJ. Capsid size and deoxyribonucleic acid length: the petite variant of bacteriophage T4. J Virol. 1970;6:865–76.

[14] Ackermann HW. Frequency of morphological phage descriptions in the year 2000. Arch Virol. 2001;146:843–57.

[15] Fokine A, Chipman PR, Leiman PG, Mesyanzhinov VV, et al. Molecular architecture of the prolate head of bacteriophage T4. Proc Nat Acad Sci. 2004;101:6003–8.

[16] Olson NH, Gingery M, Eiserling FA, Baker TS. The structure of isometric capsids of bacteriophage T4. Virology. 2001;279:385–91.

[17] Mosig G, Carnighan JR, Bibring JB, Cole R, Bock HGO, Bock S. Coordinate variation in lengths of deoxyribonucleic acid molecules and head lengths in morphological variants of bacteriophage T4. J Virol. 1972;9:857–71.

[18] Doermann AH, Eiserling FA, Boehner L. Genetic control of capsid length in bacteriophage T4 I. Isolation and preliminary description of four new mutants. J Virol. 1973;12:374–85.

[19] Cummings DJ, Chapman VA, DeLong SS, Couse NL. Structural aberrations in T-even bacteriophage. III. Induction of “lollipops” and their partial characterization. Virology. 1973;54:245–61.

[20] Keller B, Dubochet J, Adrian M, Maeder M, Wurtz M, Kellenberger E. Length and shape variants of the bacteriophage T4 head: mutations in the scaffolding core genes 68 and 22. J Virol. 1988;62:2960–9.

[21] Moody MF. The shape of the T-even bacteriophage head. Virology. 1965;26:567–76.

[22] Cummings DJ, Couse NL, Forrest GL. Structural defects of T-even bacteriophages. In: Smith KM, Lauffer MA, Bang FB, Editors. Advances in virus research. Academic Press; 1970. p.1–41

[23] Eiserling FA. T4 structure and initiation of infection, Structure of the T4 virion. In: Mathews CK, Kutter EM, Mosig G, Berget PB, Editors. Bacteriophage T4, ASM 1983, p.11–24.

[24] Mosig G. Coordinate variation in density and recombination potential in T4 phage particles produced at different times after infection. Genetics. 1963;48:1195–200.

[25] Lane T, Serwer P, Hayes SJ, Frederick, E. Quantized viral DNA packaging revealed by rotating gel electrophoresis. Virology. 1990;174:472–8.

[26] Miller ES, Kutter E, Mosig G, Arisaka F, Kunisawa T, Rüger W. Bacteriophage T4 genome. Microbiol Mol Biol Rev. 2003;67:86–156.

[27] Paulson JR, Lazaroff S, Laemmli UK. Head length determination in bacteriophage T4: the role of the core protein P22. J Mol Biol. 1976;103:155–74.

[28] Smith KO, Trousdale M. Multiple-tailed T4 bacteriophage. J Bacteriol. 1965;90:796–802.

[29] Pfau CJ, Holt SC. Multitailed T2 bacteriophage. J Virol. 1967;1:1087–8.

[30] Ackermann HW, Dauguet C, Paterson WD, Popoff M, Rouf MA, Vieu JF. *Aeromonas* bacteriophages: reexamination and classification. Ann Inst Pasteur Virol. 1985;136:175–99.

[31] Voelker R, Sulakvelidze A, Ackermann HW. Spontaneous tail length variation in a *Salmonella* myovirus. Virus Res. 2005;114:164–6.

[32] Rao VB, Black LW. Structure and assembly of bacteriophage T4 head. Virol J. 2010,7:356.

[33] Kramberger P, Honour RC, Herman RE, Smrekar F, Peterka M. Purification of the *Staphylococcus aureus* bacteriophages VDX-10 on methacrylate monoliths. J Virol Meth. 2010;166:60–4.

[34] Korol N, Van den Bossche A, Romaniuk L, Noben JP, Lavigne R, Tovkach F.. Experimental evidence for proteins constituting virion components and particle morphogenesis of bacteriophage ZF40. FEMS Microbiol Lett. 2016;363(6).

[35] Joshi A, Siddiqi JZ, Rao GR, Chakravorty M. MB78, a virulent bacteriophage of *Salmonella typhimurium*. J Virol. 1982;41:1038–43.

